# Integration of expression datasets to identify biomarkers for accurate Gleason scoring in Prostate Cancer

**DOI:** 10.1101/2024.08.21.608965

**Authors:** Pedro Matos-Filipe, Baldo Oliva, Judith Farrés, José Manuel Mas

## Abstract

Prostate cancer is a significant global health issue with considerable mortality rates, emphasising the urgent need for advanced treatment options and improved diagnostic methods. Current diagnostic standards for prostate cancer, including PSA testing and digital rectal examination, often produce false positives, resulting in unnecessary biopsies for patients. This limitation highlights the critical requirement to incorporate more precise biomarkers to enhance diagnostic accuracy and reduce unnecessary procedures. This study aims to investigate biomarker candidates that can effectively determine prostate cancer aggressiveness. By integrating diverse prostate tissue expression datasets and employing machine-learning techniques, this approach seeks to refine diagnostics and provide insights into the molecular underpinnings of the disease, potentially transforming early detection and patient management strategies. Our proposed biomarkers achieve a minimum precision of 0.80, addressing the false positives limitations associated with classical prostate cancer biomarkers. Moreover, the ROC-AUC profiles of most of the candidates proposed in this study align with those exhibited by other innovative biomarkers recently proposed (ROC-AUC ≥ 0.70). We believe these biomarkers are promising candidates for further *in vivo* and *in vitro* investigation.

## Introduction

Prostate cancer is a significant health concern, being the second most common cancer in men and the fifth leading cause of cancer-related death in males worldwide [1]. The disease can be multifactorial, involving genetic predisposition, acquired genetic mutations, and environmental influences that affect critical signalling pathways regulating the growth of cancer cells [2]. The treatment of prostate cancer has seen significant advancements, with various therapy options available, including androgen-based therapies (inhibition and deprivation) [3, 4], chemotherapy [5, 6], poly-ADP-ribose polymerase (PARP) inhibitors [7, 8], and radiopharmaceutical therapy [9, 10] (although these mostly focus on metastastatic cases). Regardless, most currently available therapeutic strategies focus on late-stage cancer cases (i.e. metastatic castration resistant prostate cancer - mCRPC) [11], while local cancer is ofter treated with through prostatectomy.

Initial detection usually occurs through the measurement of prostate-specific antigen (PSA) levels in blood, alongside Digital rectal examination (DRE) [12, 13] to assess the prostate’s size, texture, and any abnormalities. Upon suspicious findings, a biopsy may be conducted. Through this process, the presence of cancer cells is confirmed and cancer stage is determined. However, such procedure may lead to mild to severe complications, such as bleeding (i.e., haematuria, haematochezia, haematospermia), pain, urine retention, erectile dysfunction [14–16], and infection [17], which may lead to more serious complication if untreated. This makes biopsy an overbearing procedure to the patient. On the other hand, given the impact of complications, the aid of imaging techniques is usually applied prior to biopsy (i.e. MRI or CT scans). However, these are expensive procedures, which makes them more frequent in wealthier health systems [18, 19]. Furthermore, given the shared relation to non-pathological phenotypes (i.e. prostatitis and benign prostatic hyperplasia), both PSA measurements and DRE are prone to false positives [20–23] which leads to some patients being derived to biopsy without having clinically significant disease. The integration of a broader array of biomarkers, including genetic [24], epigenetic [25], transcriptomic [26], and proteomic markers [27], enables a more nuanced and precise diagnostic framework [28]. This multi-faceted biomarker approach not only enhances the accuracy of early detection but also improves patient stratification, allowing for more precise prognostication and individualized treatment planning.

Staging helps determine the aggressiveness of the tumor and the likelihood of cancer spreading. The grading system for the classification of prostate cancer tumours involves a combination of factors: (1) PSA measurements, (2) Gleason score, and (3) TNM classification [29–31]. Gleason scoring assesses the hystologic appearance of prostate cancer cells, grading them on a scale from 1 to 5 based on their differentiation and architecture [32, 33]. The total Gleason Sum (GS), derived from the two most prevalent patterns observed, helps classify the aggressiveness of the cancer in a scale of 2 to 10. Cancers classified with GS7 are special clinical cases. Gleason score of 3+4 indicates that the majority of the tumour is composed of moderately differentiated cells (grade 3) with a smaller proportion of poorly differentiated cells (grade 4), suggesting a less aggressive cancer. Conversely, a score of 4+3 indicates a higher proportion of poorly differentiated cells (grade 4) compared to moderately differentiated cells (grade 3), implying a more aggressive and potentially more dangerous cancer [34, 35]. Concurrently, the TNM staging system [36] which describes the local invasion and tumour size (T), lymph node invasion (N), and metastasis (M), evolved to provide a standardised framework for assessing cancer progression. Over the years, both systems have been refined and integrated into clinical practice, with the Gleason score serving as a vital component within the broader context of the TNM staging system [37].

Expression data is pivotal in discovering novel disease biomarkers, significantly improving patient stratification in clinical settings. By analysing complex gene expression patterns across diverse diseases and patient cohorts, researchers can identify specific molecular signatures that differentiate disease subtypes [38], predict prognosis [39], and inform treatment decisions [40]. This comprehensive molecular profiling enables the discovery of biomarkers with greater sensitivity and specificity than traditional diagnostic markers, thereby advancing disease classification and personalised therapeutic strategies. Furthermore, integrating gene expression data is a powerful approach to enhancing scientific understanding by consolidating information from multiple studies. By amalgamating data from various publications, researchers can create larger and more diverse datasets, providing comprehensive insights into biological mechanisms and disease processes [41, 42]. This integration allows the identification of subtle trends and patterns that may not be detectable in smaller, isolated datasets. Additionally, integrated datasets increase statistical power [43, 44], facilitating more robust analyses and validation of findings across different experimental conditions and populations. These insights from expression data improve our understanding of disease mechanisms and advance precise healthcare interventions, leading to better patient outcomes and the acceleration of tailored therapies.

The primary objective of this study is to identify novel biomarkers that can accurately determine the Gleason Score (GS) in prostate cancer patients. By combining microarray data from various GEO experiments using a novel method, we aim to enhance diagnostic precision by integrating these biomarkers with those currently in use. This approach seeks to provide a more comprehensive understanding of the disease’s molecular characteristics and reduce false positives and negatives, addressing a major challenge in prostate cancer diagnostics.

## Methods

### Expression dataset

An integrated dataset comprising microarray gene expression data was extracted from the Anaxomics Real-World Data repository (axRWD) [45]. The data in this repository is processed using an especial microarrays processing technique based on machine learning (ML) and the expression of anchor House-Keeping genes, called MACAROON. MACAROON allows the incorporation of microarrays samples from the Gene Expression Omnibus (GEO) repository [46] into new integrated datasets, allowing the analysis of higher sample sizes. The dataset included all samples identified with by the terms “PROSTATIC NEOPLASMS” and “PROSTATIC HYPERPLASIA”. The GEO metadata from these samples was recaptured, and samples with annotation on GS were kept. This resulted in a gene expression dataset of prostate tissue from pca patients. Given that none of the selected samples had annotation on the deconvoluted Gleason Score, all annotations were annotated using Gleason Sum (single-digit annotation). Additional clinical information (i.e., age, sex, PSA levels, sample tissue, biochemical recurrence, and clinical T stage) was mined using a tailored rule-based [47] text-mining script. Given the known clinical difference between samples classified GS 3+4 and GS 4+3, two analysis datasets were created (1) including GS=7 and (2) excluding GS=7. The final list of patient samples, and corresponding GEO series is available in supplementary table . *log*_2_ transformation was applied to expression data.

### Machine-learning models

ML models were trained with the objective of understanding which genes were better able to discriminate between samples with low-GS (GS<7) and high-GS (GS≥7). The analysis was conducted using the Gaussian Naive Bayes algorithm, as implemented in the Python package *scikit-learn* v. 1.5.0 [48]. To ensure the relevance of the ML models, genes present in fewer than 25 samples were excluded from the experiment.

Training was executed using a 10-fold cross-validation strategy, incorporating permutation tests to assess the statistical significance of the models. Models yielding non-significant p-values (*p* > 0.05) were discarded to maintain robustness. Additionally, performance was further assessed using a test set comprising a random selection of 10% of the samples, providing an extra layer of evaluation to validate the model’s generalisability.

### Selection of biomarker candidates

Predictive potential was assessed by means of accuracy (equation 1), precision (equation 2), recall (equation 3), area under the receiver-operating characteristic curve (ROC-AUC) (equation 4), and F1-metric (equation 5). Biomarker candidates were selected on the basis of cross-validation precision, for that, a cut-off of 0.8 was selected.

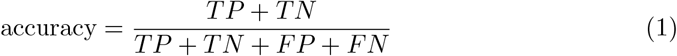

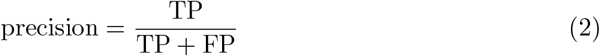

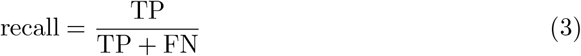

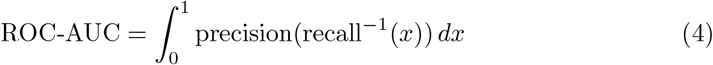

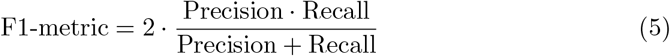

where TP = True Positives; FP = False Positives; TN = True Negatives; and FN = False Negatives.

### Differential expression analysis

In both datasets, a differential expression (DE) analysis was conducted in order to infer clinical significance of predictions. Log Fold Change (Log FC) was calculated as described in equation 6. Note that the calculation is based on subtraction given that samples were originally in the *log*_2_ form in the dataset. Statistical significance was assessed by applying the Welch’s t-test given the high occurrence of different sample sizes and variances on the two test populations [49]. Calculations were performed in Python using the package *scipy* v. 1.13.1 [50].

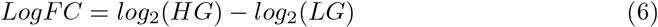

where HG = High-grade prostate cancer (GS>7; or GS≥7); LG = Low-grade prostate cancer (GS<7).

### Functional analysis

Protein-protein interactions (PPI) between the selected biomarker candidates and proteins known to be involved with prostate cancer were analysed to assess the biological significance of the selected biomarkers. Prostate cancer-related proteins were selected from the Biological Effectors Database (BED) [51]. The BED includes literature-derived proteins and genes known to be linked to medical conditions and their functional motifs. The total BED description used in this analysis, and corresponding reference, is available in supplementary table . PPI information was obtained from the STRING database [52]. Visual analysis was performed using Cytoscape v. 3.9.1 [53].

## Results

### Overview of the integrated Prostate Cancer dataset

This study focuses on the exploration of a gene expression dataset drawn from the Anaxomics’ RWD repository with the objective of finding new high-grade prostate cancer biomarkers. The repository includes all patient-derived microarrays samples in GEO as of 22/12/2020, which amounts to a total of 1,866,292 microarrays samples. For this experiment, samples with clinical tags “PROSTATIC NEOPLASMS” and “PROSTATIC HYPERPLASIA” were extracted, which resulted in 22,698 samples. In order to focus the analysis on the exploration of expression differences among patients with high-grade prostate cancer (GS≥7) against patients with low-grade prostate cancer (GS<7), we used a rule-based text-mining script to select, from the raw GEO metadata, which samples included annotation on GS. This final filter resulted in the final sample set applied in this paper, which includes a total of 870 prostate cancer samples, with expression data of 18,557 genes. The absence of annotation for the deconvoluted Gleason Score resulted on the amalgamation of the two Gleason Score “gray zone” classifications: 3+4 (considered tumour stage II – intermediate (favourable) prostate cancer) and 4+3 (consider tumour stage III – intermediate (unfavourable) prostate cancer) [54, 55]. To study the impact of this amalgamation, the global dataset was separated into three classes: (1) GS<7, (2) GS=7, and (3) GS>7; leading to the composition of two subdatasets to be used in later steps: (1) Including GS7 (where GS7 was included in the high-grade prostate cancer class), and (2) Excluding GS7. The clinicopathological description of the prostate cancer patients is presented in Table 1. The table shows that, according to available data, despite the slight increase of average age of patients with higher GS, differences are not statistically significant (Pearson correlation: *ρ* = 0.1018, *p* = 0.1211). On the other hand, other clinical variables linked to cancer severity show significant correlation with GS, specifically the average PSA level (Pearson correlation: *ρ* = 0.1155, *p* = 0.0146), biochemical recurrence (*χ*^2^ : *p* = 0.0416), and clinical stage (*χ*^2^ : *p* = 0.0003). However, the dataset originates from the integration of different gene expression datasets, with different metadata availability, result in a high proportion of missing values varies across variables. This affects the overall completeness of the dataset, especially in what concerns biochemical recurrence and sample tissue.

**Table 1.** Clinical details of the PCa individuals included in this study. GS: Gleason sum; Std. Dev.: Standard deviation; MV: Missing values; cT: clinical stage.

| Variable | Analysis Cohort |  |  |  |  |  |  |  |  | Total |  |  |
| --- | --- | --- | --- | --- | --- | --- | --- | --- | --- | --- | --- | --- |
|  | Average | *GS < 7*<br>Std. Dev. | % MV | Average | *GS = 7*<br>Std. Dev. | % MV | Average | *GS > 7*<br>Std. Dev. | % MV | Average | Std. Dev. | % MV |
| *Age* | 62 | 6.127 | 83.33% | 61 | 6.182 | 67.15% | 64 | 7.413 | 71.67% | 62 | 6.551 | 73.22% |
| *PSA (ng/mL)* | 8.74 | 5.110 | 57.97% | 11.49 | 8.927 | 45.65% | 21.03 | 53.759 | 41.67% | 13.02 | 27.357 | 48.74% |
| Variable | N | % | % MV | N | % | % MV | N | % | % MV | N | % | % MV |
| *Sample tissue* |  |  | 32.61% |  |  | 56.76% |  |  | 43.89% |  |  | 46.44% |
| Prostate | 186 | 67.39% |  | 179 | 43.24% |  | 101 | 56.11% |  | 466 | 53.56% |  |
| *Biochemical Recurrence* |  |  | 79.35% |  |  | 73.67% |  |  | 81.11% |  |  | 77.01% |
| Yes | 16 | 5.80% |  | 52 | 12.56% |  | 16 | 8.89% |  | 84 | 9.66% |  |
| No | 41 | 14.86% |  | 57 | 13.77% |  | 18 | 10.00% |  | 116 | 13.33% |  |
| *cT stage* |  |  | 5.07% |  |  | 1.93% |  |  | 9.44% |  |  | 4.48% |
| TX | 1 | 0.36% |  | 0 | 0.00% |  | 0 | 0.00% |  | 1 | 0.11% |  |
| T1 | 4 | 1.45% |  | 2 | 0.48% |  | 0 | 0.00% |  | 6 | 0.69% |  |
| T2 | 162 | 58.70% |  | 187 | 45.17% |  | 79 | 43.89% |  | 428 | 49.20% |  |
| T3 | 70 | 25.36% |  | 183 | 44.20% |  | 67 | 37.22% |  | 320 | 36.78% |  |
| T4 | 25 | 9.06% |  | 34 | 8.21% |  | 17 | 9.44% |  | 76 | 8.74% |  |
| *Gleason sum* |  |  | 0.00% |  |  | 0.00% |  |  | 0.00% |  |  | 0.00% |
| 0 | 1 | 0.36% |  | 0 | 0.00% |  | 0 | 0.00% |  | 1 | 0.11% |  |
| 4 | 3 | 1.09% |  | 0 | 0.00% |  | 0 | 0.00% |  | 3 | 0.34% |  |
| 5 | 15 | 5.43% |  | 0 | 0.00% |  | 0 | 0.00% |  | 15 | 1.72% |  |
| 6 | 257 | 93.12% |  | 0 | 0.00% |  | 0 | 0.00% |  | 257 | 29.54% |  |
| 7 | 0 | 0.00% |  | 414 | 100.00% |  | 0 | 0.00% |  | 414 | 47.59% |  |
| 8 | 0 | 0.00% |  | 0 | 0.00% |  | 113 | 62.78% |  | 113 | 12.99% |  |
| 9 | 0 | 0.00% |  | 0 | 0.00% |  | 67 | 37.22% |  | 67 | 7.70% |  |
| **Total** | 276 | 100.00% |  | 414 | 100.00% |  | 180 | 100.00% |  | 870 | 100.00% |  |

### Impact of Gleason Sum 7

Gaussian Naive Bayes models were trained in order to select genes with good predictive capability to distinguish between high-grade (GS≥7) and low-grade (GS<7) prostate prostate cancer patients. These models took as input the expression of each one of the genes in the dataset and, as output, the prostate cancer grade of the patient (High-grade, or Low-grade). The choice for this model algorithm had the objective of having the gaussian profile of *log*_2_ microarrays data into consideration, while maintaining a simple model architecture given the low number of features taken by each model. This step was performed in the two datasets: (1) Including GS7 and (2) Excluding GS7. Table 2 compares the overall performance metrics of the classification models for predicting prostate cancer severity. Supplementary tables (dataset including GS7) and (dataset excluding GS7) show all the prediction results, including test metrics. The metrics evaluated include accuracy, precision, recall, and Area Under the Curve (ROC-AUC). Excluding GS7 patients from training shows higher mean and median accuracy overall, suggesting that more genes are capable of distinguishing higher GS vs lower GS. Precision and recall are higher overall when GS7 samples are included in training, indicating that the genes are better at correctly identifying true negatives, but also true positives when these samples are taken into account. Regardless, ROC-AUC values are similar between the two groups, with slightly higher values (although with a statistically significant difference – *p* = 2.0412 × 10^−54^) when GS7 is excluded from training. This suggests comparable overall model performance in distinguishing between classes in both datasets. Where not reported, p-values in a Mann-Witney U rank test of medians were bellow python’s minimal floating point values, showing extremely high statistical significance.

**Table 2.** Global metrics for models trained with datasets including (GS7) and excluding patients with Gleason Sum 7 (No GS7). The asterisks indicate statistical significance (p¡0.05) in a Mann-Witney U rank test.

| Metric | Including GS7 | Excluding GS7 |
| --- | --- | --- |
| ***Accuracy \**** |  |  |
| Mean | 0.6148 | 0.7055 |
| Median | 0.6113 | 0.7104 |
| Std. Dev. | 0.0533 | 0.0575 |
| Min | 0.1417 | 0.3033 |
| Max | 0.8422 | 0.8633 |
| ***Precision \**** |  |  |
| Mean | 0.8536 | 0.0679 |
| Median | 0.9600 | 0.0000 |
| Std. Dev. | 0.2162 | 0.1314 |
| Min | 0.0000 | 0.0000 |
| Max | 1.0000 | 0.9633 |
| ***Recall \**** |  |  |
| Mean | 0.6331 | 0.1203 |
| Median | 0.6338 | 0.0000 |
| Std. Dev. | 0.0570 | 0.1902 |
| Min | 0.0000 | 0.0000 |
| Max | 0.8849 | 0.8000 |
| ***ROC-AUC \**** |  |  |
| Mean | 0.5648 | 0.5730 |
| Median | 0.5754 | 0.5867 |
| Std. Dev. | 0.0677 | 0.0901 |
| Min | 0.1333 | 0.0833 |
| Max | 0.8500 | 0.9500 |

### Diagnostic performance of genes for high-grade PCa

As explained in the introduction of this paper, the most relevant drawback in the most common Prostate Cancer biomarkers in clinical practice is the high False Positive rate. This supports that newly proposed biomarkers, if studied alongside current solutions, should focus on the maximisation of precision, as it measures the proportion of true positive classifications among all positive results. In other words, high precision reflects a greater proportion of true positive classifications relative to false positives. Consequently, Table 3 shows the overall performance metrics of the candidates whose models comply with a cross-validation precision cut-off of 0.8000. To guide the reader, other metrics reaching values of at least 0.7500 are marked in bold. ML results were complemented by a class permutation test [56] to account for the statistical significance of training. Models with a cross-validation accuracy p-value above 0.05 were discarded from the final list. From the initial list of 18,557 genes, these criteria allowed for the selection of 33 genes when training includes GS7, and three genes when it excludes GS7. When incorporating GS7 in the dataset, the only gene to be able to reach the 0.80 mark in precision while attaining at least 0.75 in all other assessed metrics was CCDC180. Additionally, two other genes complying with precision cut-off (ACTG1 and SERINC) were able to go beyond 0.75 in accuracy, F1-metric and recall.

**Table 3.** Selected biomarker candidates. From the 18,557 genes tested, the table shows those that were able to achieve cross-validation precision of at least 0.80 († in header) in both experiments (i.e. including patients with GS=7, and excluding patients with GS=7). Apart from ML metrics the table shows log Fold Change from differential gene expression analysis. The asterisk denotes statistical significance (*p* < 0.05).

| Uniprot ID | Gene Name | Cross-validation metrics |  |  |  |  |  |
| --- | --- | --- | --- | --- | --- | --- | --- |
|  |  | Accuracy | ROC-AUC | F1 | Precision † | Recall | Log FC |
| *Including GS=7* |  |  |  |  |  |  |  |
| P0DPF3 | NBPF9 | 0.7274 | **0.7803** | **0.7741** | **0.8849** | 0.6969 | -1.5130 * |
| Q7Z7C8 | TAF8 | 0.7200 | **0.7611** | 0.7181 | **0.8167** | 0.7000 | -0.3549 * |
| Q5VXT5 | SYPL2 | 0.7005 | **0.7596** | **0.7666** | **0.8071** | 0.7356 | -1.1123 * |
| Q8N660 | NBPF15 | 0.7138 | **0.7564** | **0.7710** | **0.8562** | 0.7052 | -1.4074 * |
| Q9BVJ7 | DUSP23 | 0.7108 | **0.7540** | **0.7765** | **0.8094** | 0.7480 | -1.7164 * |
| P19961 | AMY2B | 0.7054 | **0.7513** | **0.7674** | **0.8253** | 0.7231 | -1.4905 * |
| Q9P1Z9 | CCDC180 | **0.7667** | **0.7500** | **0.8367** | **0.8167** | **0.9000** | **0.8030** * |
| Q7Z572 | SPATA21 | 0.7005 | 0.7490 | 0.7429 | **0.8695** | 0.6567 | -1.6872 * |
| P04746 | AMY2A | 0.7115 | 0.7484 | **0.7523** | **0.8615** | 0.6779 | -1.6789 * |
| Q8NG81 | OR2M7 | 0.7306 | 0.7482 | **0.7988** | **0.8422** | **0.7688** | -2.4123 * |
| Q7RTX7 | CATSPER4 | 0.7020 | 0.7440 | **0.7547** | **0.8545** | 0.6804 | -1.3691 * |
| Q8N4P6 | LRRC71 | 0.6916 | 0.7407 | 0.7404 | **0.8048** | 0.6947 | -1.0655 * |
| Q8NAN2 | MIGA1 | 0.6884 | 0.7382 | 0.7462 | **0.8101** | 0.7014 | -1.4932 * |
| Q68G74 | LHX8 | 0.7153 | 0.7353 | **0.7739** | **0.8332** | 0.7269 | -1.2112 * |
| Q495T6 | MMEL1 | 0.6855 | 0.7341 | 0.7266 | **0.8285** | 0.6543 | -1.6362 * |
| Q8NDY8 | TMEM52 | 0.7029 | 0.7331 | 0.7407 | **0.8486** | 0.6676 | -1.7709 * |
| Q8NEQ6 | SRARP | 0.7229 | 0.7246 | **0.7872** | **0.8138** | **0.7696** | -0.9486 * |
| Q8NAP8 | ZBTB8B | 0.7063 | 0.7185 | **0.7725** | **0.8093** | 0.7443 | -1.0348 * |
| Q8IZN7 | DEFB107A, DEFB107B | 0.7050 | 0.7167 | **0.7848** | **0.8033** | **0.8333** | -0.0673 |
| Q96RW7 | HMCN1 | 0.6855 | 0.7094 | 0.7286 | **0.8208** | 0.6674 | -1.4630 * |
| Q8NB59 | SYT14 | 0.6887 | 0.7060 | 0.7257 | **0.8440** | 0.6458 | -1.9040 * |
| A6NH21 | SERINC4 | **0.7667** | 0.7000 | **0.8233** | **0.8500** | **0.8500** | **1.4549** * |
| Q8N1B3 | CCNQ | 0.6742 | 0.6910 | 0.7181 | **0.8104** | 0.6589 | **2.1524** * |
| Q8TEY5 | CREB3L4 | 0.7003 | 0.6869 | **0.7674** | **0.8027** | 0.7397 | -1.7450 * |
| Q6ZV50 | RFX8 | 0.6081 | 0.6543 | 0.3892 | **0.8217** | 0.2786 | -0.5448 * |
| Q9UMX1 | SUFU | 0.6061 | 0.6432 | 0.4511 | **0.8281** | 0.3326 | -0.6289 * |
| Q6UWF9 | FAM180A | 0.6137 | 0.6151 | 0.4864 | **0.8195** | 0.3600 | -0.4287 * |
| Q8N8Z6 | DCBLD1 | 0.5603 | 0.6057 | 0.3028 | **0.8683** | 0.1964 | -0.1249 |
| Q3SX64 | CIMAP1D | 0.5909 | 0.6022 | 0.4123 | **0.8264** | 0.2879 | -0.1606 |
| Q8TBB0 | THAP6 | 0.5961 | 0.5896 | 0.4298 | **0.8282** | 0.3061 | -0.6750 * |
| Q8IW50 | FAM219A | 0.5913 | 0.5778 | 0.4261 | **0.8229** | 0.3053 | -0.4369 * |
| Q96EY5 | MVB12A | 0.5963 | 0.5743 | 0.4290 | **0.8232** | 0.3061 | -0.4418 * |
| P63261 | ACTG1 | **0.7867** | 0.5550 | **0.8724** | **0.8083** | **0.9550** | 0.8690 * |
| *Excluding GS=7* |  |  |  |  |  |  |  |
| Q14582 | MXD4 | 0.7147 | 0.6990 | 0.5073 | **0.8000** | 0.4050 | **0.6521** * |
| O43812 | DUX1 | 0.6878 | 0.6205 | 0.3600 | **0.8000** | 0.2333 | -0.5312 * |
| O14981 | BTAF1 | 0.7372 | 0.5615 | 0.3074 | **0.8000** | 0.1976 | -0.7469 * |

Given that the proposed biomarker candidates are intended to complement those already in clinical practice (i. e., PSA), similar models to those trained using exclusively expression data were trained to include expression data of selected genes alongside the patient PSA blood level. Figure 1 shows the ROC curves for models trained with and without PSA levels as features. The complete list of results is available at supplementary table . Generally, the inclusion of PSA levels into the models’ training sets improved the quality metrics of those models. In fact, median accuracy raised to 0.8078 (maximum: OR2M7 – 0.8825), F1-metric to 0.8847 (maximum: OR2M7 – 0.9374), precision to 0.8481 (maximum: NBPF9 – 0.8914), and recall to 0.9173 (maximum: OR2M7 – 1.0000). On the other hand, ROC-AUC gets reduced (Pearson correlation: *ρ* = −0.2919, *p* = 0.1317) when gene expression data is combined with PSA levels, showing a median ROC-AUC of 0.6404 (maximum: CIMAP1D – 0.7833). However, the data shows that those models able to get a ROC-AUC increase by including PSA levels are the ones with the worst results when tested without it (i.e., FAM180A, DCBLD1, CIMAP1D, THAP6, FAM219A, and MVB12A).

**Fig 1.**
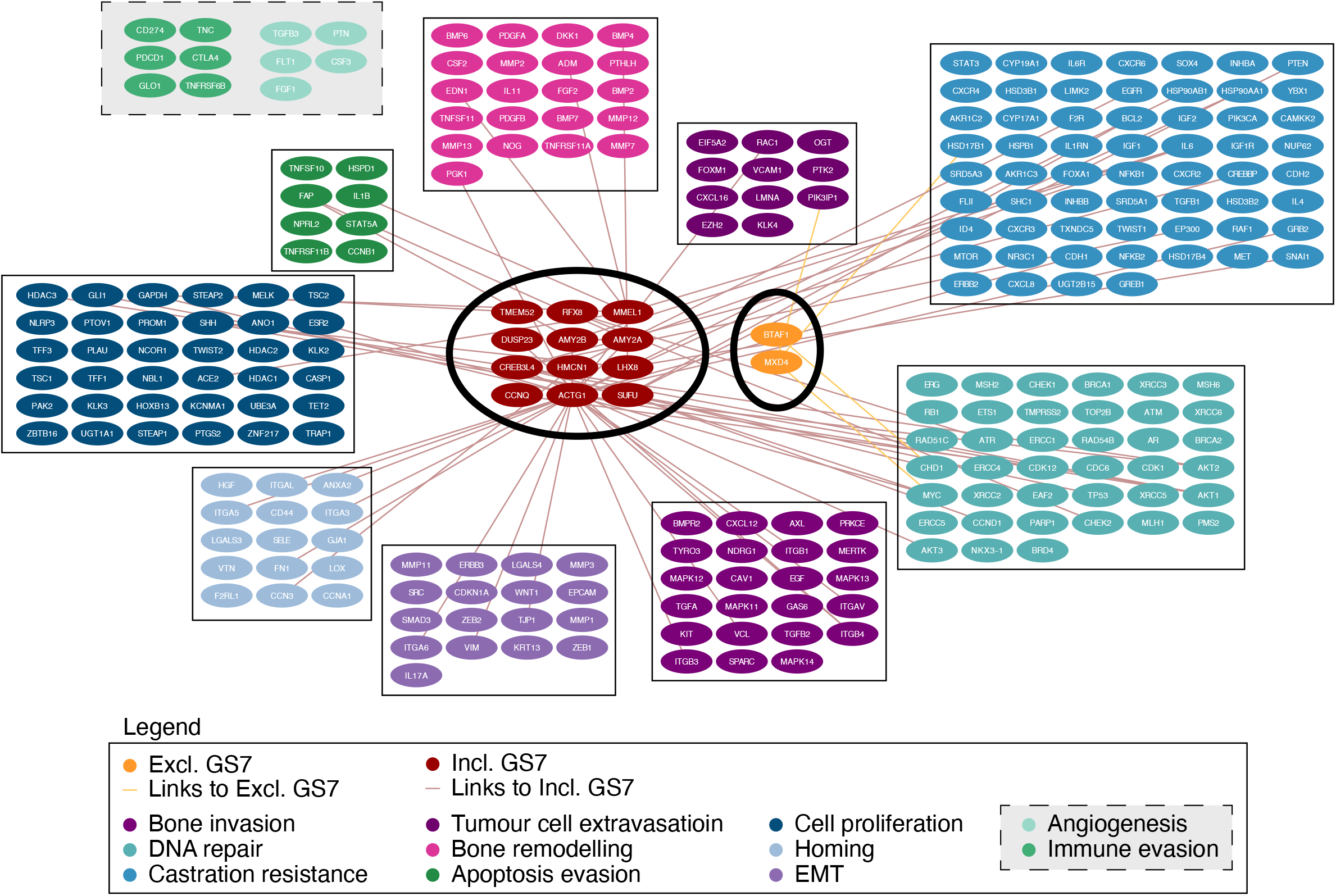
Receiver operating characteristic (ROC) curves showing the predictive efficiency of the candidate biomarkers for distinguishing high-grade from low-grade Prostate Cancer patients. In each subplot, we present the ROC curves for the corresponding gene individually, PSA individually and the two biomarkers together. In cases where the combination curve is not present, the number of patients with known PSA and expression level of said gene where bellow the sample size cut-off.

From the clinical practice perspective, apart from their presence in readily accessible biological material (e. g., urine, saliva, or blood), the most useful biomarkers are usually those who are more upregulated as disease progresses. The DE profile of the genes available in the dataset was studied as means of aiding in the selection of the best suited candidates. In Table 3, we show those expression patterns for the genes complying with the ML-metric cut-offs. Extended results are available in supplementary table and supplementary figure . The analysis was performed separately on both prostate cancer datasets. This resulted in the identification of 36 differentially expressed genes including GS7 samples in the dataset (five upregulated in high-grade prostate cancer patients, and 31 downregulated), and 159 genes by excluding GS7 samples (61 upregulated, and 98 downregulated). From these lists, 28 common gene expression patterns were identified in both datasets. Four of these genes were upregulated (i. e., ZNF737, CCNQ, TENT5B, SPRYD4). However, only one (CCNQ) of these four genes was classified as candidate biomarker by the ML-based experiment. The remaining 24 were downregulated, including 7 that were considered as potential biomarker candidates by ML models: AMY2A, DUSP23, TMEM52, CREB3L4, SPATA21, OR2M7, SYT14.

### Functional analysis of identified candidates

A functional analysis was performed to understand the role of selected biomarker candidates in the context of Prostate Cancer, and to validate these genes in the context of the aetiology of the disease. To achieve this goal, the disease description of Prostate Cancer from the BED was queried alongside the selected biomarker candidates in the STRING database. This provided the PPI links between biomarker candidates that complied with ML cut-offs and the various biological biological mechanisms that are comprised in the BED’s description of Prostate Cancer, specifically: (1) Bone invasion and metastatic growth, (2) DNA Damage Repair (DDR) Alterations, (3) Castration resistance, (4) Tumour cell extravasation, (5) Pathologic bone remodelling, (6) Evading apoptosis, (7) Cell growth and Proliferation, (8) Cancer cell homing in metastasis, (9) Epithelial to mesenchymal transition (EMT), (10) Sustained Angiogenesis, (11) Immune system evasion. A representation of the aforementioned PPIs is displayed in Figure 2. From the 36 selected biomarker candidates, two were not mapped to any protein in STRING (i. e., CIMAP1D, and DUX1). From the remaining 34, 14 had known direct PPIs with proteins in the BED molecular description of Prostate Cancer. By biological mechanism: (1) Bone Invasion: DUSP is linked with bone invasion through its interaction with ITGB4, highlighting its role in the metastatic spread of cancer cells to bone tissue. (2) Apoptosis Evasion: AMY2B and AMY2A are associated with apoptosis evasion via FAP, with AMY2B also connected to ACTG1, another candidate biomarker. MMEL1 is linked to apoptosis evasion through IL1B. (3) Castration Resistance: AMY2B and AMY2A are linked to castration resistance via IL6. LHX8 is involved in castration resistance through SNAI1, and MMEL1 through IL6. CREB3L4 is associated with castration resistance through interactions with CREBBP, EP300, and AKT2. SUFU is connected to castration resistance via BCL2 and PTEN, while MXD4 is linked through HSD17B1. (3) Cell Proliferation: LHX8 is related to cell proliferation via SHH, MMEL1 via ACE2 and GAPDH, and TMEM52 through STEAP2. CREB3L4 is linked to cell proliferation through HDAC3, while SUFU connects via CCND1, GLI1, and SHH. ACTG1 shows strong connections to cell proliferation through ESR2 and GAPDH. (4) Bone Remodelling: LHX8 is connected to bone remodelling via BMP4, and MMEL1 through ADM and EDN1. ACTG1 is linked to bone remodelling via PGK1. (5) Homing: HMCN1 is associated with homing via CCN3, and ACTG1 through ITGAL, ITGA5, and ITGA3. (6): Epithelial-Mesenchymal Transition (EMT): HMCN1 is linked to EMT via EGFR, while ACTG1 is connected through FN1, TJP1, ITGA6, VIM, CDH1, and ITGB4. MXD4 is also involved in tumour cell extravasation through PIK3IP1. Lastly, (7) DNA Damage Repair: CCNQ is related to DNA damage repair through CDK12, CREB3L4 through AKT1, AKT3, and CHD1, RFX8 via ATR and CHEK2, SUFU through TP53, AKT1, and MYC, ACTG1 via MYC, and MXD4 through MYC. BTAF1 is connected to DNA repair through CHD1.

**Fig 2.**
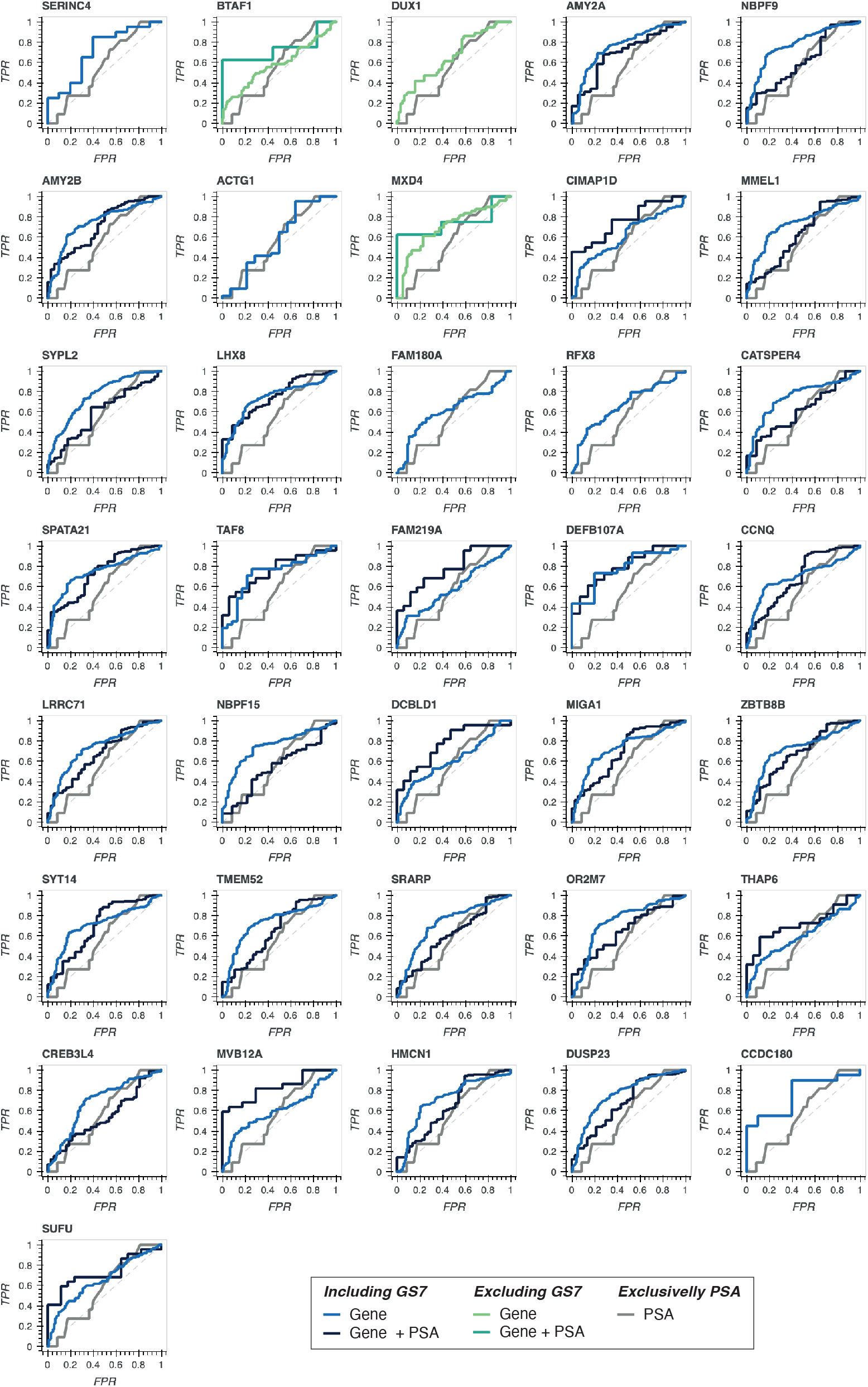
Known protein-protein interactions between selected candidates and key Prostate Cancer motifs. Identified candidates are marked by the oval shapes in the centre of the image. The legend of the figure follows the same order as the protein clusters represented, and should be read in the anti-clockwise sense. GS: Gleason Sum; EMT: Epithelial to mesenchymal transition.

## Discussion

Among the first line diagnostics for Prostate Cancer patients is the evaluation of blood PSA levels, and DRE. However, the low specificity and sensitivity of current diagnostics, specifically in what concerns PSA, is a well recognised problem that often leads to unnecessary biopsy. In this study, we present a list of promising new biomarker candidates that show potential for the reduction of false positives through the maximisation of precision. To achieve this objective, we have resorted to a gene expression dataset obtained by integrating data from different GEO experiments. The genes included in this dataset were scored by training ML models (i.e., Gaussian Naïve Bayes classifiers) where the expression of each gene was matched to a prostate cancer class: (1) low-grade if GS<7, or (2) high-grade if GS≥7. Given that information on the decomposed GS was unavailable (separate Gleason scores at the two prostate regions), we tested the impact of having GS7 included in the dataset, despite comprising patients with different clinical outcomes (3+4 – favourable, and 4+3 – unfavourable). Our results show no relevant differences in the global classification metrics obtained using either dataset. This, although somewhat surprising, is common in other proposed biomarkers (discussed below). Another issue that arose due to insufficient metadata was the inability to analyse the effect of the prostate sample’s origin in the patients included in the datasets, as this information was missing and could not be incorporated into the analysis. Cancer heterogeneity within the same prostate gland is a known factor with practical consequences on the final GS attributed to the patient. Knowing if GS was the sum of two Gleason scores obtained from biopsies of radial, imaging-guided, or prostatectomy-derived biopsies may have substantially impacted on the global quality and composition of the dataset. On the other hand, the microarrays data present in the dataset was normalised using a process (MACAROON) that uses global values across the whole axRWD repository as anchor points. This implies that the expression values in the dataset do not directly correspond to those presented by patients. Consequently, the models adjusted to this dataset may not be directly applicable to patient-derived data. Instead, numerical values would need to be subjected to the same transformations as the data included in the axRWD repository, which may be impractical in a clinical setting. For this reason, the biomarker candidates presented in this work serve to shed light onto new unexplored molecules and to support further study both *in vitro* as well as in patients.

Other recently discovered (some approved in selected markets) biomarkers show improvement compared to PSA at identifying clinically significant prostate cancer (GS ≥ 7). Two frequently explored biological sample sources are blood and urine. Blood biomarkers include the 4K panel, which includes the assessment of protein levels of total PSA, free PSA, intact PSA, and human kallikrein 2 (hK2) [57] obtaining a GS≥7 ROC-AUC of 0.78 [58]. Other approved biomarker panel is the Prostate Health Index (PHI), which includes the protein level of free PSA, total PSA, and the [-2] form of proPSA (p2PSA), reaching an ROC-AUC of 0.72 [59, 60]. Urine biomarkers often resort to mRNA readings. Successful examples include PCA3 (ROC-AUC=0.703 [61]) Select MDx (HOXC6, DLX1; ROC-AUC 0.83 [62]); ExoDx Prostate (ERG; ROC-AUC=0.74 [63]). The Mi-Prostate Score uses sequencing information to assess T2-ERG fusion alongside expression of PCA3, and serum PSA, reaching an ROC-AUC of 0.77 [64]. In our *in silico* experiments, presented candidates show comparable ROC-AUC levels, which positions them as viable experimental options. Furthermore, despite having the origin of our evidence in mRNA expression from prostate tissue, proteins corresponding to some of the presented candidates have been previously identified in extracellular vesicles (EVs). AMY2B was identified in human urine [65] and in EVs released by prostate cancer cells [66]. AMY2A was identified in human urine [65] and saliva [67], and in EVs released by prostate cancer cells [66]. MVB12A was identified in human urine [68, 69], and in EVs released by prostate cancer cells [66] (among other cancer types [70, 71]). ACTG1 was identified in human urine [65, 68, 72], and in EVs released by prostate cancer cells [73] (among other cancer types [74, 75]).

Biomarkers in current clinical practice are able to achieve such high levels ROC-AUC by building their classifiers with further clinical and demographic information; specifically, PSA levels, age, life-style, ethnical background, family history, among others [76, 77]. We attempted to perform such analysis by including PSA levels as features of the classification models which was able to enhance the predictive capacity of some candidates (i. e., FAM180A, DCBLD1, CIMAP1D, THAP6, FAM219A, and MVB12A). However, our approach based on the integration of public microarrays data implied a penalty on the availability of such information in most clinical/demographical features (often completely lacking or missing in more than 50% of subjects in most genes). On the other hand, in the case of PSA, other ML model architectures may lead to better classification metrics since PSA levels do not assume a gaussian distribution [78]. We believe that, by combining such information, the biomarker candidates proposed on this publication would achieve higher predictive scores and allow more personalised treatment strategies. Furthermore, combining the candidates identified in this paper into a new biomarker panel could also contribute to increased predictive capacity, enhancing overall diagnostic and prognostic accuracy.

In conclusion, our study integrates gene expression data from various GEO experiments and applies machine learning models to identify promising new biomarker candidates for prostate cancer diagnostics. These candidates demonstrate potential for reducing false positives and enhancing diagnostic precision. A key contribution of our work is the exploration of a large, integrated patient dataset, which significantly improves statistical power. Additionally, biomarker research in Prostate Cancer typically either targets early disease detection or more advanced stages (e.g., metastatic castration resistant prostate cancer). Our study focuses on the evolution of the Gleason Score, an aspect often overlooked in biomarker research for prostate cancer. Despite the challenges posed by insufficient metadata to conduct further testing, our results indicate that these candidates perform comparably to existing biomarkers in terms of ROC-AUC levels. Moreover, the presence of some candidates in extracellular vesicles underscores their potential applicability in non-invasive diagnostics. While the inclusion of clinical and demographic information could further refine predictive capabilities, our findings provide a valuable foundation for future studies and the development of more personalised treatment strategies in prostate cancer management.

## Supporting information

**S1 Fig. Differentially expressed genes on High GS when comparing to Low GS**. (a) corresponds to the dataset that includes GS=7; while (b) corresponds to the dataset excluding GS=7. Upregulated and downregulated genes are represented in green and red, respectively. The names of genes identified in both datasets are annotated in the plots. The dashed lines correspond to the fold-change (LogFC – i.e., -1.5 and +1.5) and p-value (*−log*_10_(*p − value*) – 0.05) cut-offs.

**S1 Table. List of GEO samples comprised in the Prostate Cancer patient dataset**.

**S2 Table. BED description of Prostate Cancer and corresponding motifs. S3 Table. Global ML prediction results (dataset including GS7)**.

**S4 Table. Global ML prediction results (dataset excluding GS7)**.

**S5 Table. ML metrics from models trained with gene expression and PSA blood levels**.

**S6 Table. ML metrics from models trained with gene expression and PSA blood levels**.

## Acknowledgments

The authors would like to thank Dr. Eva Coppola for their contribution on reviewing this manuscript.

## Availability of data and materials

Code is available on github

(github.com/ppintofilipe/gleason-scoring-biomarkers). Datasets and supplementary data were made available on Zenodo (zenodo.org/records/13120584).

## Authors’ contributions

PMF: methodology, software, validation, formal analysis, investigation, Writing – original draft, Writing – review and editing, visualisation. BO: Supervision, Resources, Writing – review and editing. JF: Conceptualisation, Funding acquisition, Investigation, Project administration, Supervision, Validation, Writing – review and editing. JMM: Methodology, Resources, Supervision, Writing – review and editing.

