## Supplementary figures and images for "Integration of expression datasets to identify biomarkers for accurate Gleason scoring in Prostate Cancer"

### Supplementary Figure 1

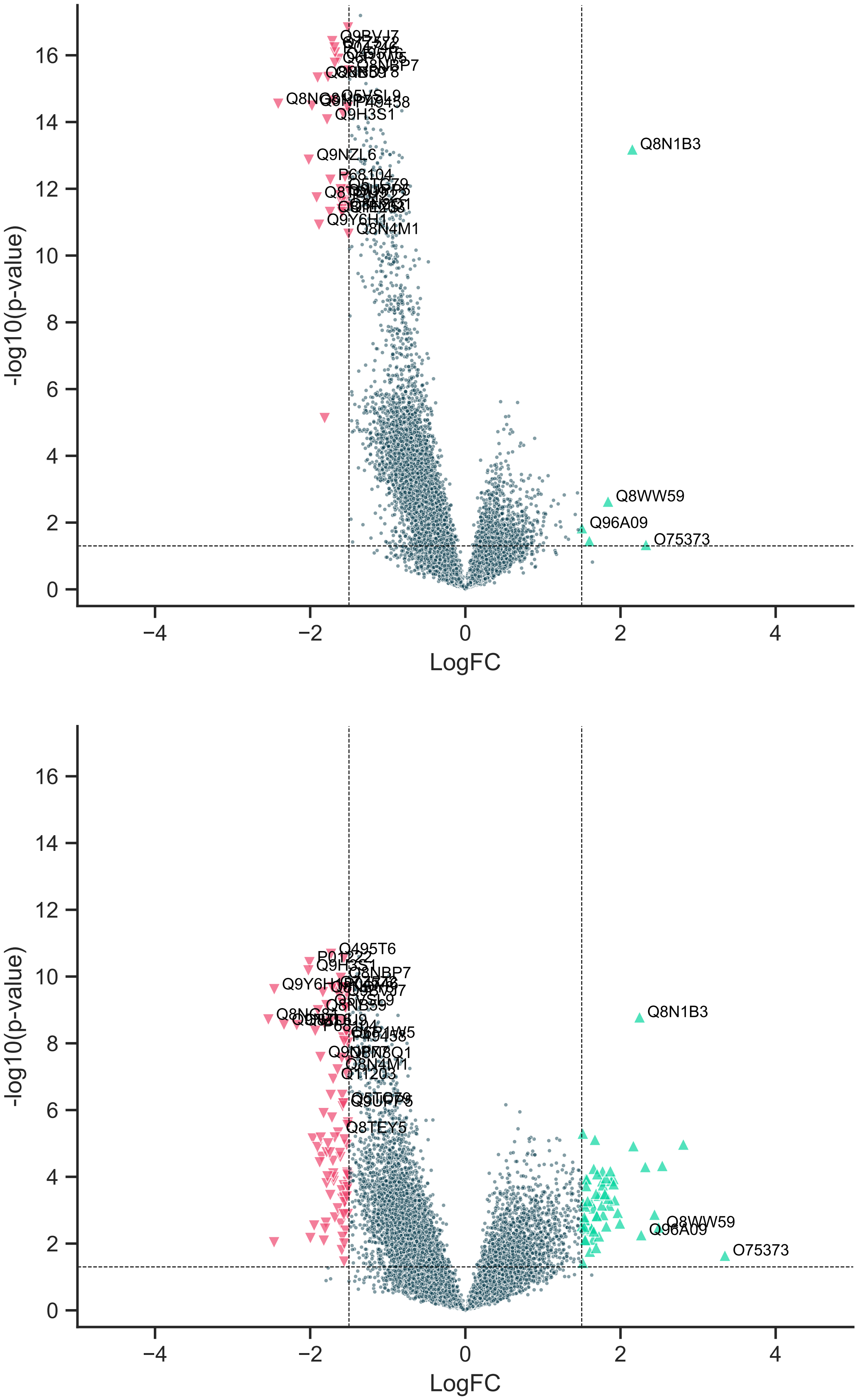
